# Phenotypic consequences of logarithmic signaling in MAPK stress response

**DOI:** 10.1101/2023.12.05.570188

**Authors:** Hossein Jashnsaz, Gregor Neuert

## Abstract

How cells respond to dynamic environmental changes is crucial for understanding fundamental biological processes and cell physiology. In this study, we developed an experimental and quantitative analytical framework to explore how dynamic stress gradients that change over time regulate cellular volume, signaling activation, and growth phenotypes. Our findings reveal that gradual stress conditions substantially enhance cell growth compared to conventional acute stress. This growth advantage correlates with a minimal reduction in cell volume dependent on the dynamic of stress. We explain the growth phenotype with our finding of a logarithmic signal transduction mechanism in the yeast Mitogen-Activated Protein Kinase (MAPK) osmotic stress response pathway. These insights into the interplay between gradual environments, cell volume change, dynamic cell signaling, and growth, advance our understanding of fundamental cellular processes in gradual stress environments.

## INTRODUCTION

Signal transduction is fundamental for a variety of cellular processes, including growth, differentiation, migration, and programmed cell death (1). These complex pathways play a pivotal role in maintaining normal cellular functions, and disruptions in their regulation can result in developmental defects and various human diseases (2–8). Despite the wealth of knowledge on individual proteins and signaling pathways, accurately predicting how cells respond to a range of perturbations, such as environmental changes and drug treatments, remains a formidable challenge (9–11). Understanding the mechanisms of signal transduction is a crucial step in characterizing cellular responses in both healthy and diseased states.

The complexity of cell signaling behavior is further compounded by their spatio-temporal dynamics. Current cell biology primarily focuses on studying signaling behavior by measuring a limited number of pathway components under steady-state conditions or in response to acute, step-like sudden shifts between two constant concentrations (12–14). However, in their physiological context, cells encounter stress concentrations that change in various forms such as a sudden, linear, nonlinear, and repetitive forms over time, space, or both (15, 16). These changes occur at time scales and length scales that are different from those involving acute transitions between two constant levels (17). For example, circadian oscillations occur once per day, synaptic firing happens multiple times per second, and various hormones, including blood insulin, exhibit unique temporal patterns that selectively regulate intracellular processes based on their dynamics (18). These physiological changes that cells experience are not always acute and they could happen in smooth and slow patterns over time or space (17). This mismatch between traditional experimental conditions and the dynamic reality of physiological environments has created a gap in our understanding of how signaling behaves within complex natural settings.

Emerging studies demonstrate that different cell stimulations that vary over time or space can dramatically affect intracellular signaling dynamics (17, 19). As a result, qualitatively distinct dynamics of the same signaling molecule such as signals with different amplitudes, durations, rates, or oscillations can lead to altered gene expression or create entirely distinct cell phenotypes (7, 15, 16, 18–32). Such cell stimulations vary widely in their nature from extracellular cues such as growth factors (33) or environmental stressors (22, 34, 35) to intercellular molecules including morphogens (7, 26, 36), cytokines (37–41), hormones (18), or neurotransmitters (42) and extending to intracellular signals such as DNA damage (43) or cell cycle factors (24). The observation that different concentration patterns of the same stimulation molecule over time or space create distinct cellular responses highlights a significant gap in our understanding of the underlying signaling mechanisms in gradual environments that represent physiology.

To mimic physiological cellular environments, we have developed a novel method for precisely controlling the microenvironment of individual cells by subjecting them to stress with concentrations that change gradually over time (15, 16, 27, 29). Our studies have demonstrated that gradual cell stress yield superior results compared to conventional approaches when constructing predictive mathematical models of cell signaling (15, 16). Furthermore, we and other pioneering studies have investigated the impact of varying the rate of osmotic stress over time on signaling activation and cellular survival (17, 19, 23, 26–29, 44–46). However, our understanding of how cells integrate dynamic signal features continuedly over time, such as the rate of change in stress, along with the stress concentration itself, remains unknown. In addition, how gradual stress impact cellular morphology and growth over time remains a significant knowledge gap.

In this study, we delved into the differential effects of acute stress compared to gradual physiological stress on the regulation of signaling and cell growth. Moreover, we sought to gain insights into how cells integrate gradual environmental changes into their responses over time. To address these questions, we present an integrated experimental and quantitative analytical framework designed to explore dynamic volume change and signaling processes under gradual environments. We employed the conserved Mitogen-Activated Protein Kinase (MAPK) High Osmolarity Glycerol (HOG) stress signaling pathway in the yeast *Saccharomyces cerevisiae* as a model system (47, 48). Here, we report our discovery that rate sensing and logarithmic signaling together contribute to the eukaryotic MAPK signal transduction, shedding light on how cells process dynamic stress into their responses over time.

The Hog1 pathway, among the well-studied eukaryotic signaling pathways, stands out as a blueprint for studying signaling mechanisms (16, 47). Remarkably, this pathway displays conservation in many of its constituent proteins, extending from yeast to humans. As a major MAPK signaling pathway, its fundamental role in the cellular response to osmotic stress makes it an ideal model for exploring cellular responses to gradual environmental changes over time. Numerous key studies have shed light on the significance of the Hog1 pathway in shaping yeast growth phenotypes during osmotic stress (Figures 1A) (48–50). This pathway has been the subject of extensive investigation, with researchers exploring various facets of cellular responses to osmotic stress and the central role Hog1 signaling plays in these intricate processes (29, 35, 48–52). For example, it is shown that cell volume reduction in severe osmotic stress conditions slows down Hog1 signal transduction from cytoplasm to nucleus due to molecular crowding and this has implications for a range of cellular processes including cell growth (53–56).

**Figure 1.**
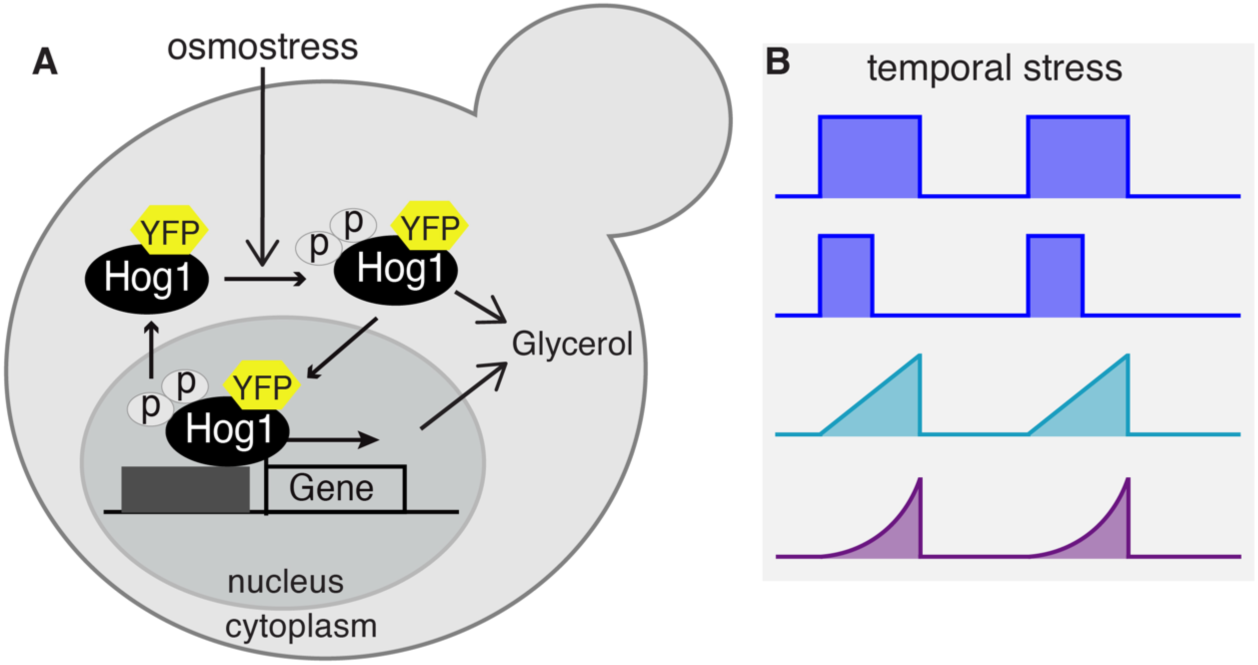
Dissecting Osmostress regulation of Volume Change, Cell Growth and Hog1 Signaling. This figure presents an illustration of the conventional Hog1 signaling in yeast stress response pathway and our proposed gradual stress paradigms. (**A**) In accordance with the established model of osmotic stress regulation in yeast, that is based on acute stress, osmotic stress activates Hog1 signaling and inhibits cell growth in an osmotic concentration stress dependent manner. Osmotic stress triggers the activation of and double phosphorylation of Hog1 MAPK, via the HOG signal transduction pathway. Dual Hog1 phosphorylation leads to its translocation to the nucleus, where it regulates stress-responsive genes. The concerted action of Hog1 phosphorylation and the gene responses contributes to the regulation of glycerol production and maintenance, facilitating osmoadaptation. (**B**) We applied different stress paradigms including acute (blue) and gradual (cyan and purple) changes to investigate the relationship between osmotic stress, cell volume change, Hog1 signaling, and cell growth.

In experiments, we employed precisely controlled gradual cell stress (Figure 1B). Through real-time measurements, we tracked cell volume changes, Hog1 nuclear localization, and cell growth phenotypes, as elaborated in the subsequent sections. Notably, we observed an optimal cellular growth phenotype under severe osmotic stresses when applied to cells gradually. Our data has revealed the existence of logarithmic signaling, a signal transduction mechanism that perceives relative changes in extracellular stress, shedding light on this phenomenon in eukaryotes with crucial implications for cell volume, signaling and growth.

In contrast to the transient Hog1 signaling activation seen with acute stress, we observed a gradual Hog1 signaling decay with linear stress and a sustained Hog1 signaling activation over the duration of exponential stress. This observation is different than the predictions of signaling mechanisms through which cells sense the concentration change or the absolute rate of change in stress, as we discuss in the results. These findings collectively contribute to our understanding of how cells adapt and respond to gradual environmental changes.

## RESULTS

### 1. Gradually Rising Stress Conditions Compared to Acute and Pulsatile Stress Conditions result in Growth Advantage

Here, we study how does gradual stress impact the cellular growth phenotype. Our findings revealed a strong correlation between the stress gradient type and cell growth. Standard yeast stress response models suggest that the severity (concentration or duration) of stress directly influences the cell growth phenotype (22, 53, 54, 56). We exposed cells to an acute stress in the form of a 3M pulsatile NaCl for 120 minutes and subsequently quantified the cell doubling time after exposure as 453+/-46 minutes (Figure 2A-2B, Methods). In comparison, exposure of cells to acute 3M NaCl stress for 30 minutes resulted in a cell doubling time of 218+/- 18 minutes (Figure 2C-2D, Methods). Next, we exposed cells to a linear increasing stress (3M NaCl in 120 minutes), and measured a reduced cell doubling time of 147+/-16 minutes (Figure 2E). Exposing cells to an exponential increasing stress scenario of the same intensity and duration reduced the cell doubling time to 117+/-1 minutes (Figure 2F). Our investigation unveiled a remarkable deviation from the standard paradigm. The type of gradual stress proved to be a pivotal factor in determining the cell growth phenotype even more than stress intensity, duration, or total stress (integrated stress exposure quantified as the total area under the stress profile) (Figure 2G-2I). Our results indicate that cells grow substantially better under gradually rising stress conditions compared to acute pulsatile stress conditions (Figure 2, purple a cyan compared to blue). These observations are even more prevalent in repetitive stress conditions after exposure to a second stress treatment (Figures 2 and 1S). This intriguing outcome sheds new light on the intricate interplay between the gradual aspects of stress and cellular responses, highlighting the significance of gradual stress exposure in enhancing cell growth.

**Figure 2.**
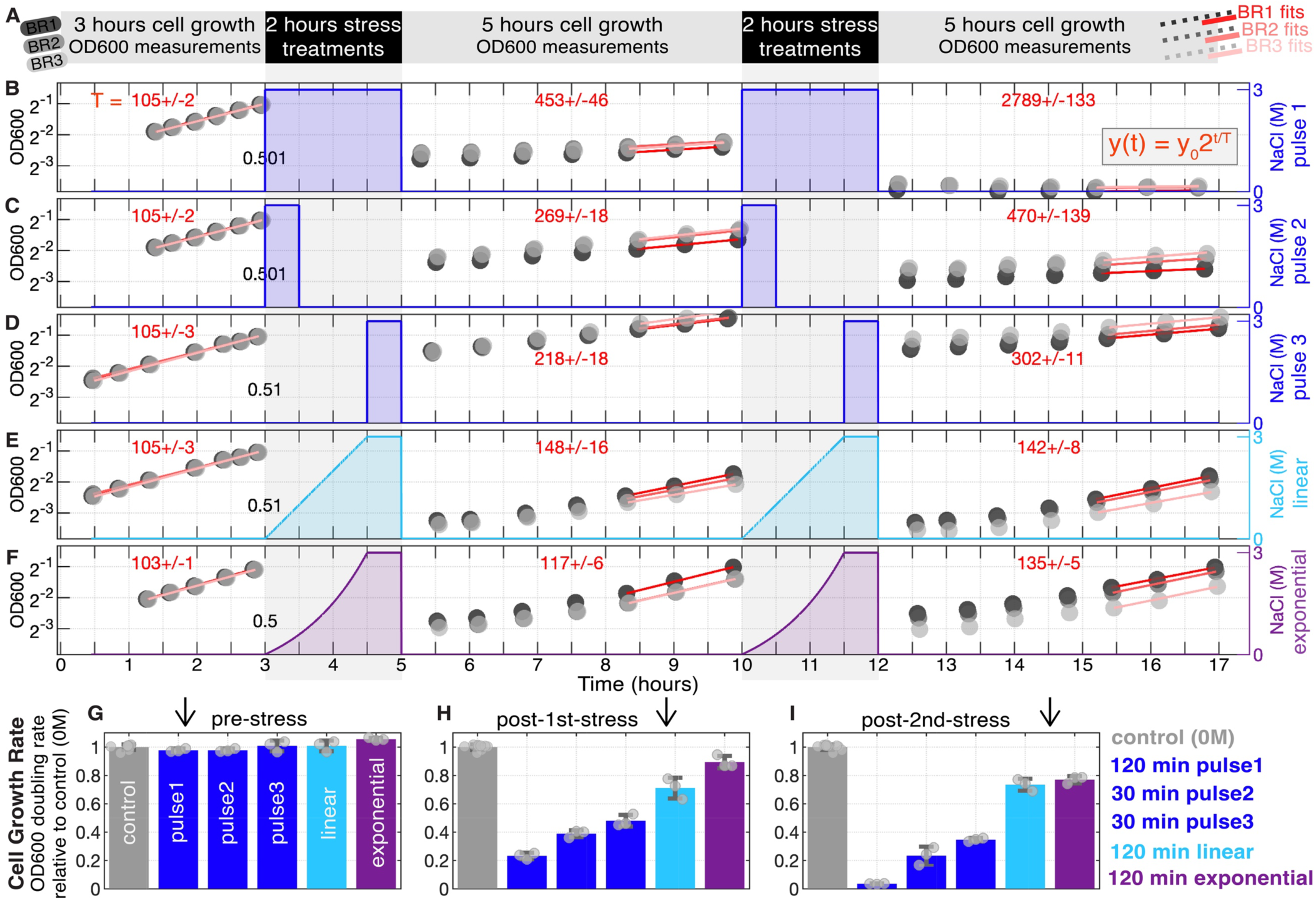
Enhanced Cell Growth under Gradual Osmotic Stress Compared to Acute Pulsatile Stress. (**A-F**) We evaluated cell growth under various NaCl stress conditions through OD600 measurements of yeast cell cultures. (**A**) Cells were initially grown to an exponential phase (OD600 = 0.5) before being subjected to repetitive 3M NaCl osmotic stresses, including (**B**) a 120-minute long pulse, (**C**) a 30-minute pulse, (**D**) a 30-minute pulse starting at t=90 min from the start of treatment period (a control for cell density, see Methods), (**E**) a 120-minute linear profile (cyan), and (**F**) a 120-minute exponential profile (purple). The linear and exponential stress profiles gradually increased to 3M over 90 minutes, remaining at this concentration for an additional 30 minutes. Following each 2-hour stress treatment, cells were filtered to remove NaCl, resuspended in fresh growth media, and their OD600 was monitored for 5 hours in 3 biological replicates (black, gray, light-filled circles in **B**-**F**). After 5 hours, the stress treatments and OD600 measurements were repeated. The OD600 measurements were fitted to exponential growth curves (**B-F**), depicted on a log2 y-axis in red over time, and fits utilized the last three timepoints. Cell growth doubling times were calculated from each fit (indicated by red numbers rounded to the nearest minute). The values provided at t=3 hours at the onset of the initial 2-hour stress treatment in (**B**-**F**) represent OD600 estimates derived from fits applied to pre-stress OD600 measurements. An approximate value of 0.5 indicates that all conditions maintained a similar initial cell density at the beginning of the 2-hour stress treatment period. (**G-I**) Cell growth rates, relative to the control (0M, no stress), were determined for (**G**) pre-stress, (**H**) post 1st stress treatments, and (**I**) post 2nd stress treatments based on OD600 doubling rates. Further details can be found in Figure S1.

### 2. Gradual Stress of Single Cells Results in Distinct Cell Volume Phenotypes and Hog1 Signaling Dynamic Responses

To understand the observed cell growth phenotypes, we implemented gradual stress paradigms and quantified cell volume changes and Hog1 signaling activation via time lapse microscopy. We aimed to elucidate these dynamics by employing precisely controlled gradual concentration profiles, contrasting them with acute stress implemented as instant step-like concentration changes. The gradual concentration profiles, implemented as polynomial functions of time (Figure 3A), were characterized by three key parameters: C_max_ (maximum concentration change), T (stress duration), and k (polynomial order), which governed how the profile reached C_max_ over T (Figure 3B). In these profiles, k = 1 represented linear stress, while k values between zero and one signified stress starting with the highest rate and gradually decreasing (Figure 3A). For k > 1, stress started with slower rates and increased over time. Additionally, we generated exponentially increasing profiles akin to the polynomial ones, beginning at 0 and reaching C_max_ over T (Figure 3B).

**Figure 3.**
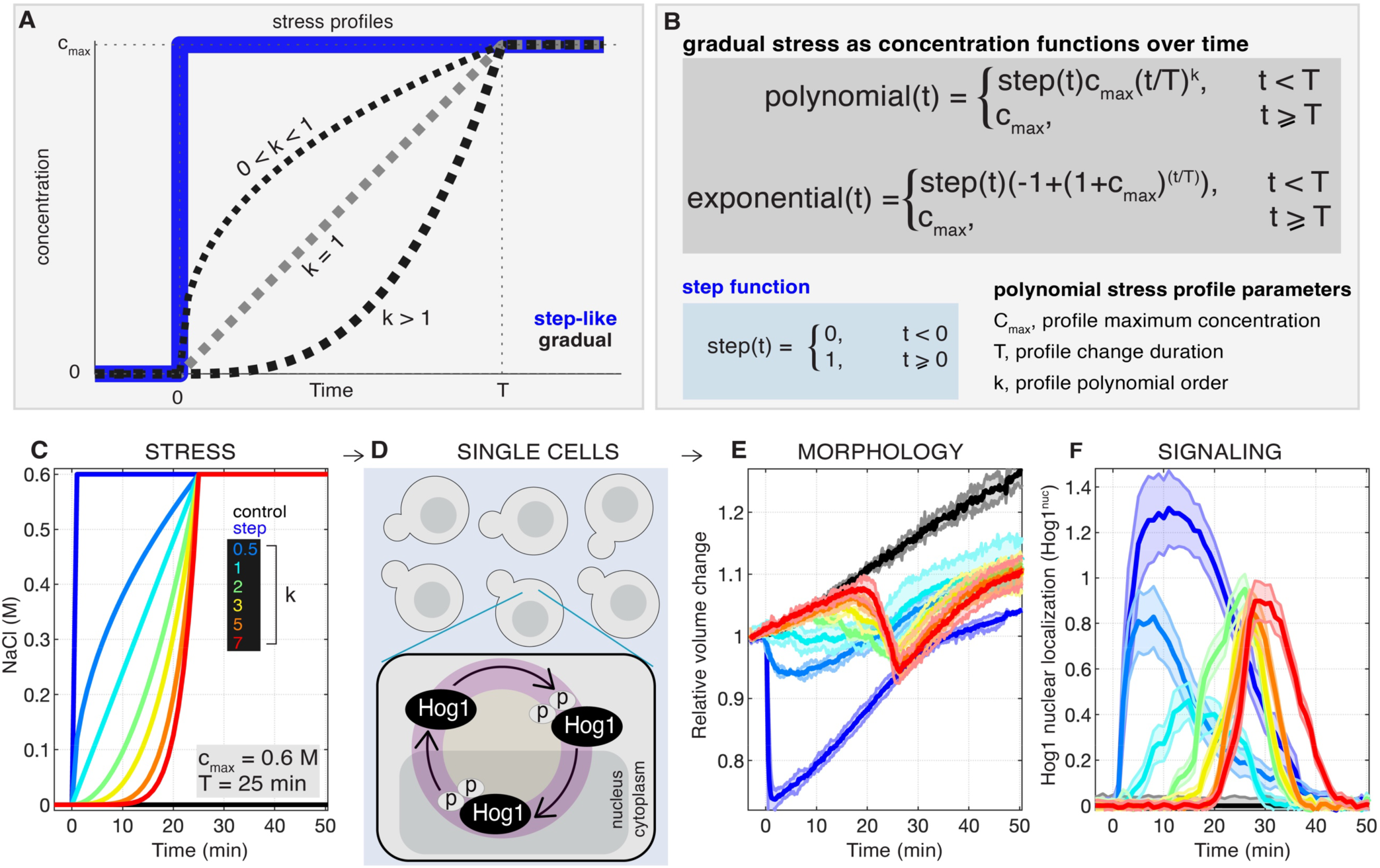
Distinct Responses to Gradual Stress in Single Cells. (**A**) We employed gradual stress concentration increases over time, contrasting them with acute, step-like stress at time zero. (**B**) Gradual stress followed polynomial increases over time, with profiles reaching the final concentration (C_max_) over a specified duration (T), determined by the polynomial order (k). Exponential increases were also considered. (**C**) Gradual NaCl stress profiles according to polynomial increases are applied to (**D**) yeast cells resulted in distinct (**E**) cell volume relative changes and (**F**) Hog1 kinase nuclear localization dynamics (Hog1^nuc^) phenotypes. Gradual NaCl concentration profiles in (**C**) followed various functions, including step (*t*^0^), root (√*t*), linear (*t*^1^), quadratic (*t*^2^), cubic (*t*^5^), quintic (*t*^5^), and heptic (*t*^7^), each reaching a final concentration of 0.60 NaCl over 25 minutes, all compared to a no-salt control (0M). The thick lines and shaded areas in (**E, F**) represent the mean and standard deviation (std) from 2 or 3 biological replicates, each consisting of approximately 100 single cells observed through live cell time-lapse microscopy. Further details can be found in Figure S2.

In our experimental setup, we compared the responses to a 0M NaCl control and an acute 0.6M NaCl stress at t=0 minutes with the responses to polynomial NaCl increases from zero to 0.6M over 25 minutes for k values of 0.5, 1, 2, 3, 5, and 7 (Figure 3C). These concentrations were then maintained at 0.6M for an additional 25 minutes (Figure 3C). Subsequently, we applied these stress profiles to single cells (Figure 3D) and conducted real-time measurements to quantify cellular volume changes over time (Figure 3E) and Hog1 nuclear localization (Figure 3F) during the stress periods using time-lapse microscopy (Figure S2).

Our results unveiled significant distinctions in cellular responses to these dynamic stress profiles even though all concentrations reached the same final values. Under growth media with 0M NaCl, cellular volume showed gradual expansion and an overall increase over time (Figure 3E). In contrast, an acute NaCl stress led to a rapid and substantial reduction in cellular volume. Gradual increases in stress resulted in gradual and sustained cell volume reductions over time (Figure 3E).

At the signaling level, we observed distinct patterns of Hog1 nuclear localization (defined as Hog1^nuc^) in response to these stress profiles (Figure 3F). In growth media with 0M NaCl, there was no discernible change in Hog1 nuclear localization. Conversely, an acute NaCl stress induced a rapid and robust transient increase in Hog1 nuclear localization. Gradual NaCl stress, however, elicited delayed Hog1 activity that did not reach the maximum activation observed under an acute stress. Furthermore, the timing of maximum activation was shifted, and all activities ultimately adapted to pre-stress levels over a 50-minute period.

Both in terms of cellular morphology and signaling dynamics, responses to gradual stress exhibited marked differences between conditions over time. These findings underscore the nuanced and distinct cellular responses arising from variations in gradual stress profiles.

### 3. Cell Volume Reduction and Hog1 nuclear localization Follow the Change in Stress Dynamics with High Fidelity

Here, we focused on quantifying the precision and fidelity of cellular responses following the gradual stress profiles, particularly their impact on cell volume reduction and Hog1 nuclear localization. Under gradual NaCl stress, with final concentrations set at C_max_ = 0.6M (Figure 4A), we observed distinct patterns in relative cell volume change (Figure 4B) and cell volume reduction (Figure 4C) phenotypes. We define cell volume reduction as the volume change under each NaCl stress condition relative to the volume change in growth media of 0M NaCl condition.

**Figure 4.**
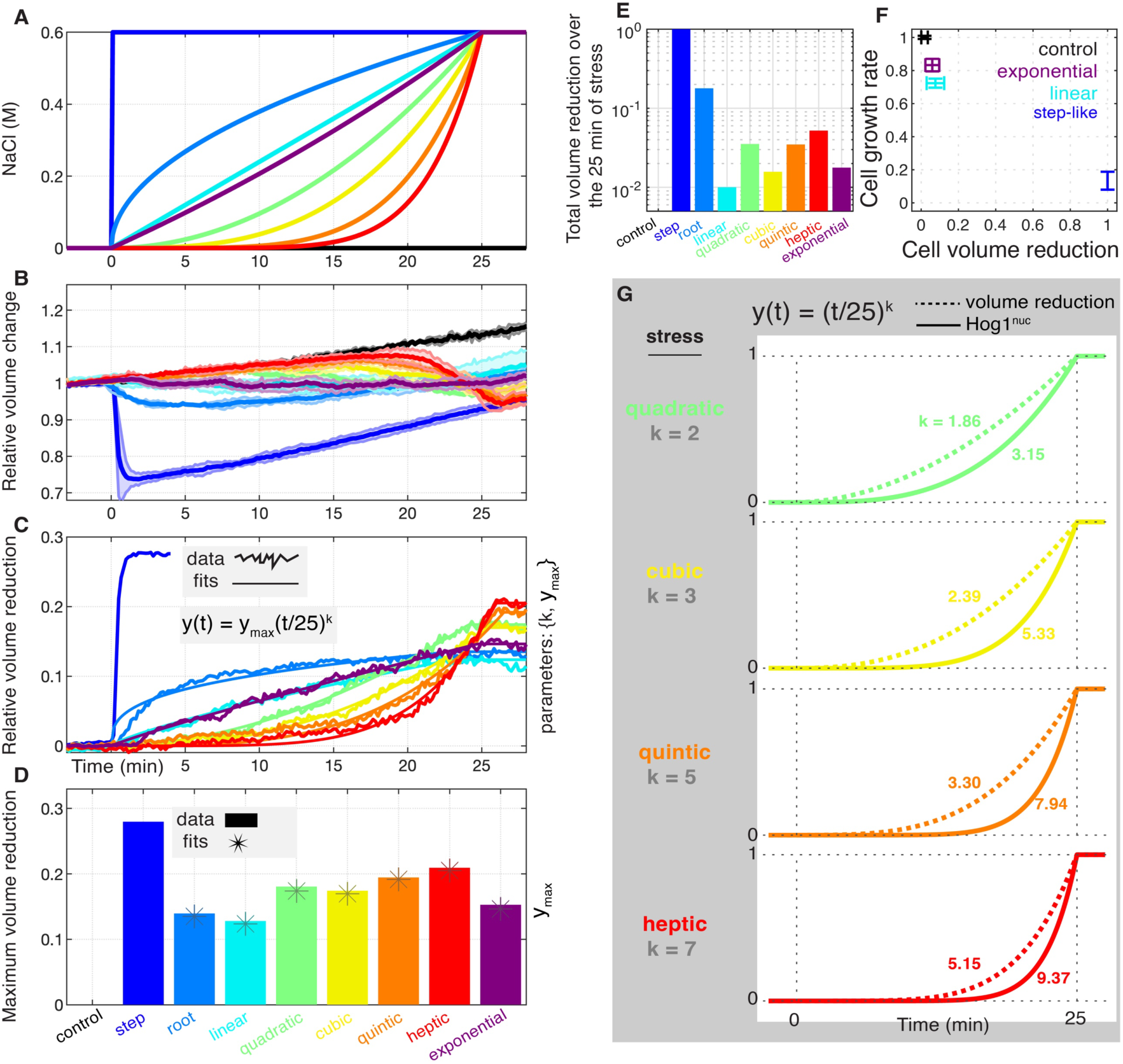
Dynamic Cellular Responses to Gradual NaCl Stress. (**A**) Gradual NaCl stress were applied to yeast cells, with final concentration of C_max_ = 0.6M, resulting in differential (**B**) relative cell volume changes and (**C**) cell volume reduction, defined as the volume change under each stress condition relative to that under the no-salt condition. (**D**) The maximum cell volume reduction and (**E**) the total cell volume reduction over the 25-minute stress period exhibited variability across different gradual stresses. (**F**) Cell growth showed a negative correlation with cell volume reduction amount, including both maximum and total reduction. The x-axis displays the mean and standard deviation (std) of volume reduction values (both maximum and total reductions) under control, step, linear, and exponential stress for five different final concentrations (C_max_ = 0.1M, 0.2M, 0.4M, 0.6M, and 0.8M), each normalized to the step condition with the greatest cell volume reduction (see Figure S3E). The y-axis illustrates the mean and std of cell growth rates under control, step, linear, and exponential stress conditions, normalized to the control condition with the highest cell growth rate values. Values from three biological replicates at post-1^st^-stress and post-2^nd^-stress are aggregated for this quantification (see Figures 2G-2I). (**G**) The dynamics of Hog1 nuclear localization (Hog1^nuc^) were delayed concerning cell volume reduction dynamics under all stress changes. Further details can be found in Figures S3 and S4.

The maximum cell volume reduction (Figure 4D) and the total cell volume reduction over the 25-minute stress time (Figure 4E) exhibited marked variations among the stress conditions. The former represents the highest reduction in cell volume, while the latter denotes the integrated cell volume reduction throughout the 25 minutes of cell stress. Notably, step-like stress resulted in the highest cell volume reductions among all the gradual stress profiles. In contrast, both linear and exponential stress led to the lowest (total or maximum) cell volume reduction when compared to other gradual stress. Furthermore, our analysis revealed a negative correlation between cell growth rate and cell volume reductions under osmotic stress conditions (Figure 4F).

Next, we quantified the dynamics of both cell volume reduction and Hog1nuc compared to NaCl stress change by fitting each volume and Hog1 response to a polynomial function (Figures 4G, S3, S4, Table 1, Methods). We found that cells process the stress change at volume level different than that of Hog1 activation. For example, in response to a quadratic stress change over time (k=2), cell volume reduction follows a polynomial change of smaller order (k=1.86) while Hog1nuc follow a polynomial change of greater order (k=3.15) (Figure 4G). This observation suggests a specific mathematical relationship between the applied gradual stress and the resulting cell volume changes and Hog1nuc dynamics.

**Table 1.**
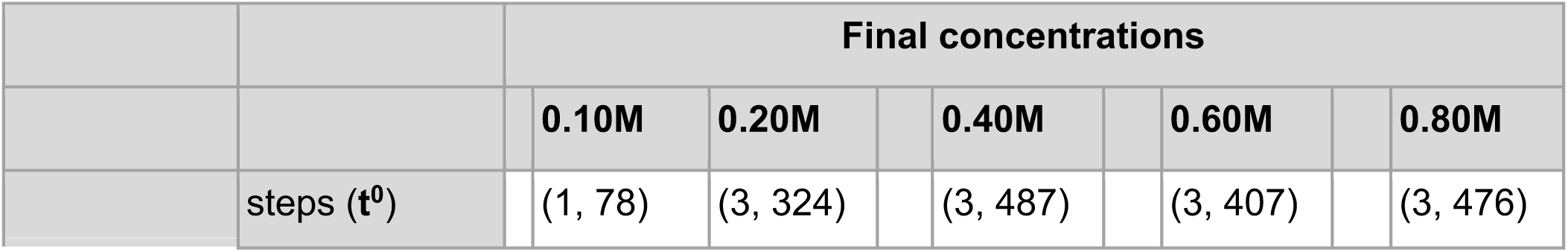
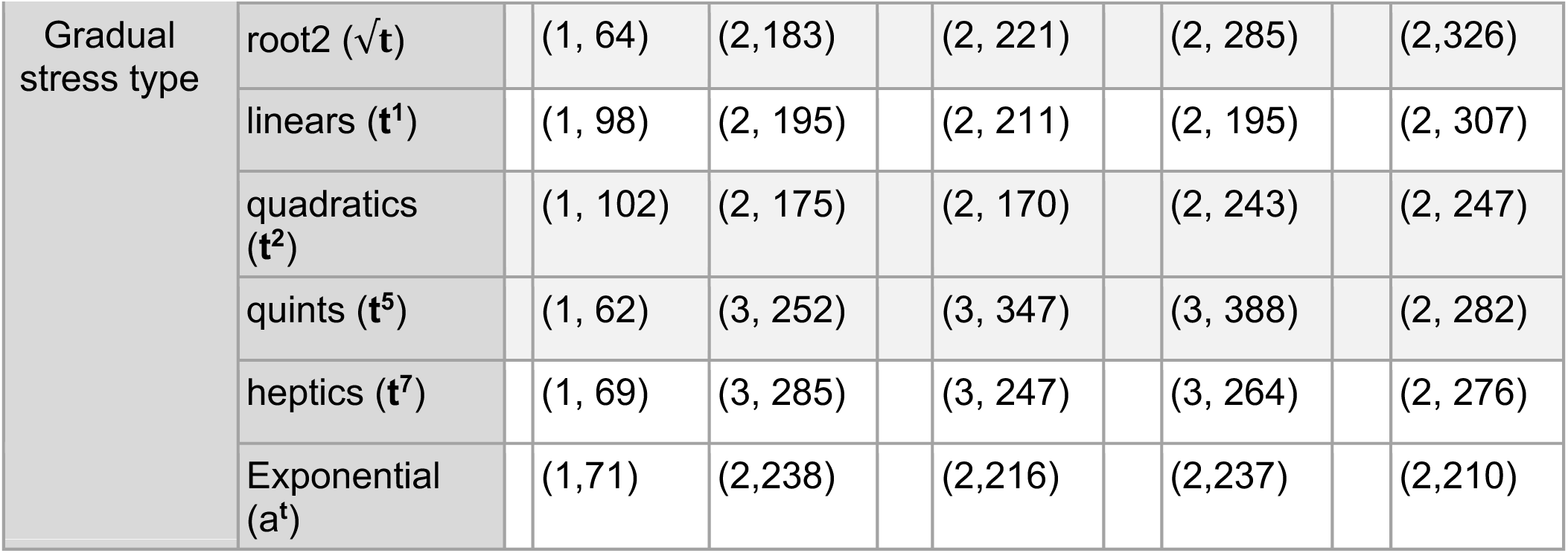
Statistics of the biological replicates and single cells used in Figures S3 and S4. For each condition, the first number represent the number of biological replicates, and the second number indicate an approximate number of total single cells from all biological replicates for that condition.

|  |  | Final concentrations |  |  |  |  |  |  |  |  |
| --- | --- | --- | --- | --- | --- | --- | --- | --- | --- | --- |
|  |  |  | 0.10M | 0.20M |  | 0.40M |  | 0.60M |  | 0.80M |
| | steps ($t^0$) | | (1, 78) | (3, 324) | | (3, 487) | | (3, 407) | | (3, 476) |

|  |  |  |  |  |  |  |
| --- | --- | --- | --- | --- | --- | --- |
| Gradual stress type | root2 ($\sqrt{t}$) | (1, 64) | (2,183) | (2, 221) | (2, 285) | (2,326) |
| | linears ($t^1$) | (1, 98) | (2, 195) | (2, 211) | (2, 195) | (2, 307) |
| | quadratics ($t^2$) | (1, 102) | (2, 175) | (2, 170) | (2, 243) | (2, 247) |
| | quints ($t^5$) | (1, 62) | (3, 252) | (3, 347) | (3, 388) | (2, 282) |
| | heptics ($t^7$) | (1, 69) | (3, 285) | (3, 247) | (3, 264) | (2, 276) |
| | Exponential ($a^t$) | (1,71) | (2,238) | (2,216) | (2,237) | (2,210) |

To answer whether Hog1nuc activity impacts cell volume reduction, we directly compared the dynamics of Hog1^nuc^ to that of cellular volume reduction under gradual NaCl stress (Figure 4G). Like acute stress conditions (23), we observed that Hog1^nuc^ dynamics is delayed relative to cell volume reduction dynamics under all gradual stress conditions. This observed delay in Hog1^nuc^ dynamics suggest that Hog1^nuc^ dynamics does not impact cell volume reduction. Rather, the observed cell volume reduction is a direct consequence of extracellular stress change, irrespective of Hog1^nuc^ activity.

These results collectively emphasize the fidelity of cellular responses to gradual stress changes, offering valuable insights into the relationship and the temporal order among extracellular stress change, cell volume dynamics, signaling activation, and cell growth phenotypes. This raises the question why do the linear and exponential profiles cause minimal volume change and enhanced growth under stress.

### 4. Logarithmic Signaling Mediates a Response Proportional to the Relative Changes of Extracellular Environment

To address this question, we aim to better understand how gradual stress profiles control Hog1 signaling dynamics by means of phenomenological predictive modeling with the aim to identify a signaling mechanism. We conducted a comprehensive analysis that compared the behaviors of linear signaling (or rate sensing) models, where the cellular response is directly proportional to the rate of change in the stress, with logarithmic signaling models, in which the response is proportionate to the relative change in stress with respect to the perceiving background concentration over time. Additionally, we explored a hybrid model that incorporated equal contributions from both linear and logarithmic signaling responses additively (see Methods). To dissect the unique characteristics of these signaling mechanisms, we subjected each of the linear, logarithmic, and hybrid models to four distinct cell stress paradigms (Figure 5).

**Figure 5.**
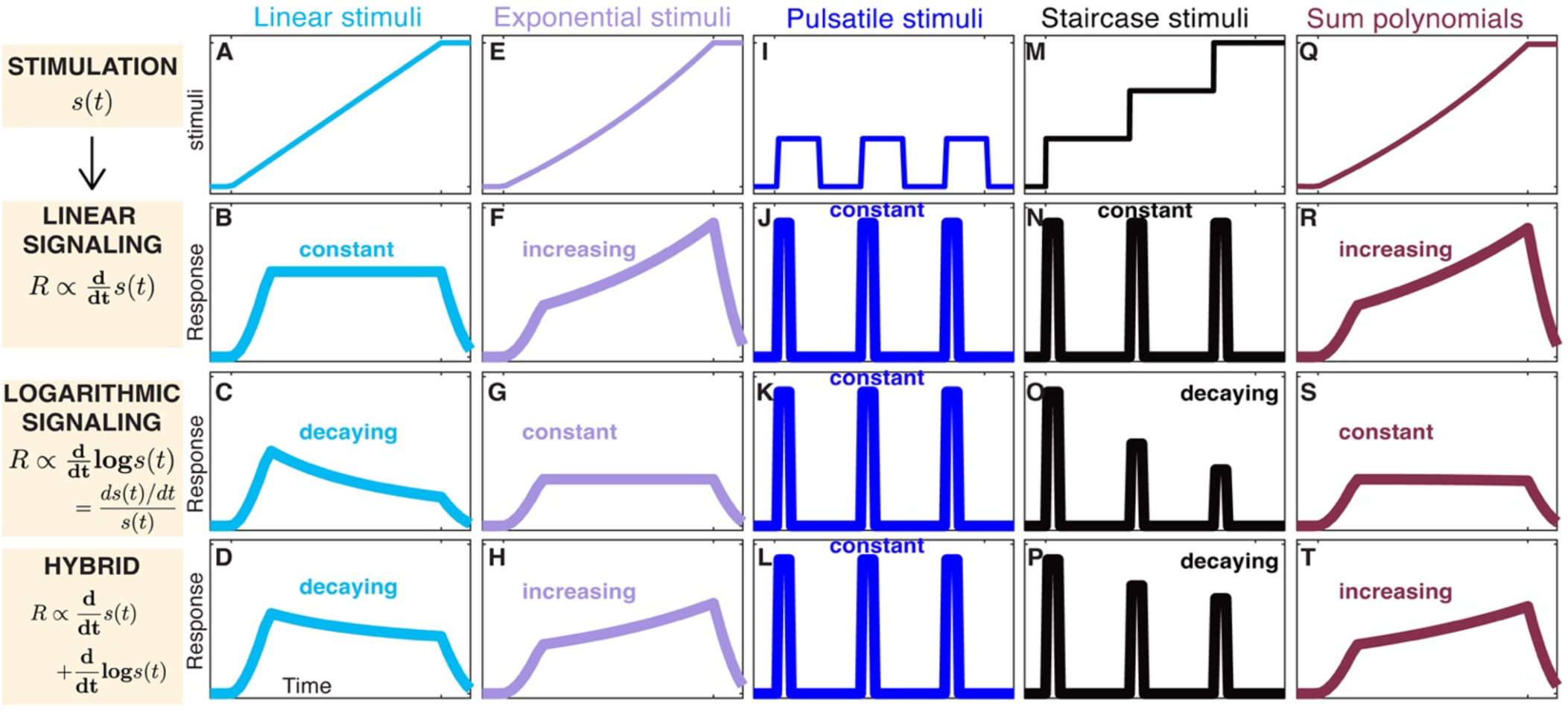
Different signaling models convert gradual environmental changes into different signaling responses. (**A**) Under a linear stress, the three models produce (**B**) constant, (**C**) decaying, and (**D**) slower decaying responses over time within the linear, logarithmic, and hybrid models, respectively. (**E**) An exponential stress results in (**F**) increasing, (**G**) constant, and (**H**) slower increasing responses over time within the three models, respectively. (**I**) Pulsatile stress induces (**J-L**) responses that remain constant throughout the stress periods across all three signaling models. (**M**) A staircase stress mediates (**N**) constant, (**O**) decaying, and (**P**) slower decaying responses throughout the stress periods within the three models. The Taylor series summation of polynomials stress of order k=1,2,3,5,7 (the Maclaurin series expansion for exponential stress, see Methods) is illustrated in (**Q**). Similar summations of signaling to polynomial stress generate (**R-T**) responses consistent with those observed with exponential stress. The responses are smoothed over 3 timepoints for each model. Further details can be found in the Methods section.

In the linear stress paradigm, cells encountered a stress that exhibited linearly increasing concentration at a consistent rate over time (Figure 5A). Our analysis predicted distinct responses under each of the three signaling mechanisms studied. Following reaching the maximum level, we predicted that the response would exhibit constant (Figure 5B), gradual decay (Figure 5C), and slower decay (Figure 5D) characteristics over time within the linear, logarithmic, and hybrid models, respectively. It’s worth noting that the hybrid model was expected to display a slower rate of decay in response compared to the logarithmic mechanism, while the linear signaling mechanism to maintain a constant response level.

In the exponential stress paradigm, cells were exposed to stress that exponentially increased in concentration over time (Figure 5E), resulting in response dynamics as follows: after the initial rise, the response was expected to show increasing, constant, and slower increasing patterns over time within the linear (Figure 5F), logarithmic (Figure 5G), and hybrid (Figure 5H) models, respectively. In each of the three signaling mechanisms, the response to an exponentially increasing stress was projected to persist at a higher level compared to the response to a linearly increasing stress.

In the pulsatile stress paradigm, cells received repeating acute stress (Figure 5I), leading to identical responses throughout the stress periods (constant responses at each pulse of stress application) across all signaling mechanisms (Figure 5J-5L). Notably, pulsatile stress did not yield distinctive responses among the signaling mechanisms, indicating their inability to discern between them.

In the staircase stress paradigm, the cells experienced a stress that followed a staircase pattern, with a constant step change at each period without switching back to pre-stress level (Figure 5M). Staircase stress differentiated between the three signaling mechanisms. The linear signaling is predicted to lead to cellular responses that remained constant throughout the stress periods (Figure 5N). On the other hand, the logarithmic and the hybrid signaling mechanisms both are predicted to lead to cellular responses that decayed throughout the stress periods (Figure 5O, 5P).

In addition to these four major cell stress paradigms and to further test different signaling paradigms, we also considered polynomial cell stress similar to those in Figure 3C. The Taylor series summation of polynomials stress profiles of order k=1,2,3,5,7 (the Maclaurin series expansion for an exponential function with coefficients as 1/k!, where k! represents the factorial of k) recapitulates an exponentially increasing stress profile (Figure 5Q compared to Figure 5E). We quantified the responses to each polynomial stress through each of the models. Similar summations of signaling responses to polynomial stress recapitulated responses consistent with those observed with exponential stress (Figure 5R-5T compared to Figures 5F-5H). Next, we experimentally validate these predictions to differentiate between linear, logarithmic, and hybrid signaling mechanisms.

### 5. Implementing Gradual Cell Stress Confirmed that Logarithmic Signaling Contributes to the HOG Pathway Signaling Response

In our experimental investigations, we employed various gradual cell stress conditions to illuminate the signaling behavior of the HOG pathway. Specifically, we examined the response of yeast cells to different forms of gradual NaCl stress at varying final concentrations. A summary of key findings are as follows:

Linear stress, characterized by a gradual increase in NaCl concentration with a constant rate over time to two different final concentrations of C_max_ = 0.6M and 0.8M (Figure 6A), induced a pattern of increasing cell volume reduction over time (Figure 6B). Quantification of these results indicates that cell volume change follows a polynomial function of order less than k=1 for the linear stress (Figures S3D-S3F, cyan). In parallel, this linear stress paradigm resulted in decaying Hog1^nuc^ dynamics after reaching the maximum level (Figure 6C, decaying trend are indicated via black lines). Exponential stress, where the NaCl concentration increased exponentially reaching two different final concentrations of C_max_ = 0.6M and 0.8M each over 25 minutes (Figure 6D), led to a linear increase in cell volume reductions (Figures 6E, S3D-S3F, purple), and an increase in Hog1^nuc^ activity over the course of stress (Figure 6F). These stress paradigms together show that the Hog1 pathway behaves as a hybrid model that combines both rate sensing and logarithmic sensing mechanisms (Figures 5D, 5H).

**Figure 6.**
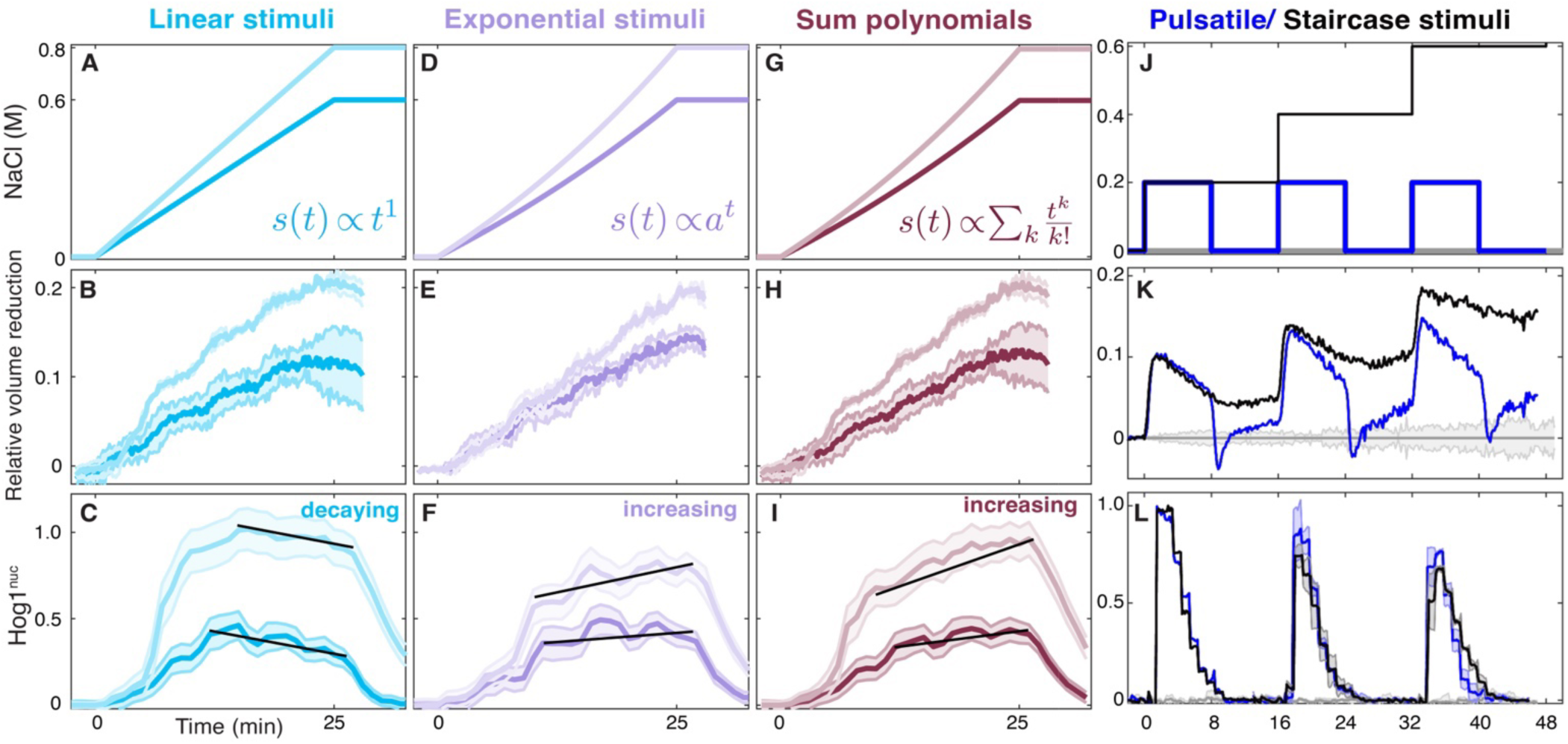
Verification of Logarithmic Signaling in the HOG Pathway through Gradual Cell Stress. (**A**) Linear NaCl stress (with final concentrations of C_max_ = 0.6M and 0.8M) lead to (**B**) increasing cell volume reduction and (**C**) decaying dynamics of Hog1 nuclear localization over time. (**D**) Exponential stress result in (**E**) increasing cell volume reduction and (**F**) increasing Hog1nuc dynamics over time. Fitting cell volume reduction to a polynomial function indicates that volume reduction changes with a smaller polynomial order under linear stress compared to exponential stress (see Figures S3D, S3F for C_max_ = 0.6M and 0.8M). (**G-I**) Summation of cell volume reduction dynamics and the transient responses of Hog1nuc to NaCl stress with polynomial orders (k=1,2,3,5,7) generated (**H**) increasing cell volume reduction and (**I**) increasing Hog1nuc dynamics over time, similar to the responses observed with exponential stress at each final concentration. (**J**) A comparison between pulsatile and staircase NaCl stress, consisting of three equal-height step elevations, shows differential dynamics in both (**K**) cell volume reduction and (**L**) Hog1nuc responses over time. The lines represent means, and the shaded areas indicate standard deviations (std) from 2-3 biological replicates (BRs), each with approximately 100 single cells analyzed through time-lapse microscopy. Further details can be found in Figures S5.

**Figure 7.**
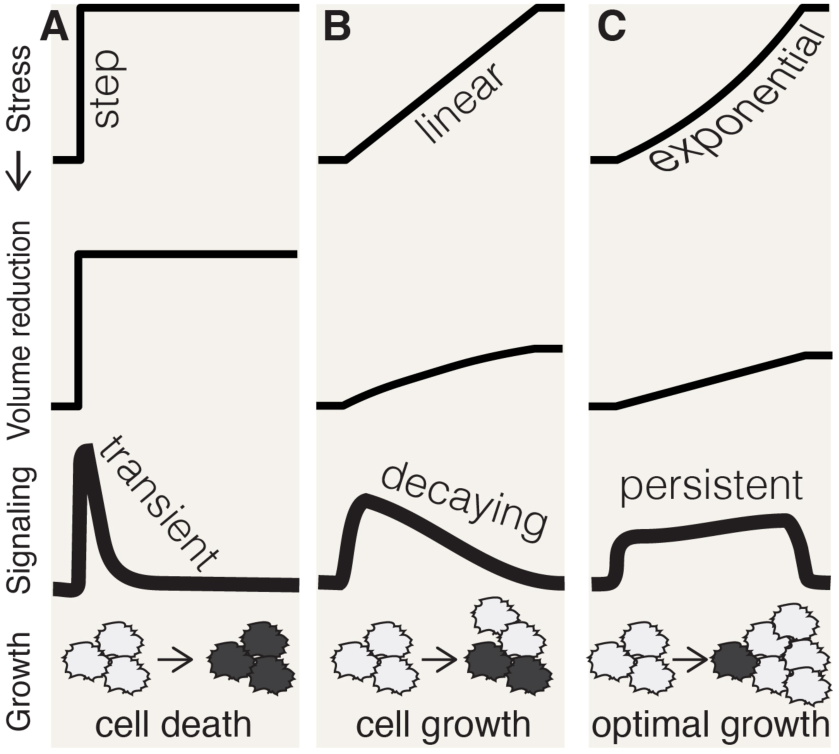
Gradual Stress Identified Logarithmic Signaling as a Mechanism Contributing to Optimal Cell Growth. (**A**) Application of different cell stress profiles over time, in the form of (**A**) an acute step change, (**B**) a linear, and (**C**) an exponential stress, yield distinctive dynamics in signaling activation dynamics and phenotypes. Notably, (**A**) rapid step stress induces substantial cell volume reduction compared to the (**B**, **C**) gradual stress, while signaling persists longer with gradual stress compared to step stress. Compared to the standard paradigm in which the concentration of osmotic stress dictates cell growth, in this study we discovered that gradual stress, in a dynamic-dependent manner, influences cell volume reduction and signaling activation, which in turn dictates the cell growth phenotype.

To further explore the response patterns, we quantified the sum of the transient Hog1^nuc^ responses induced by polynomial stress of various orders (k=1,2,3,5,7) (Figure 6G). We performed these sums using weight coefficients according to the Maclaurin series expansion (see Methods). Surprisingly, these responses closely resembled those observed under exponential stress, characterized by linearly increasing cell volume reduction and increasing Hog1^nuc^ dynamics over time (Figure 6H, 6I).

Finally, we examined the effects of two distinct stress types: pulsatile and staircase. Pulsatile stress consisted of three identical acute stresses between 0 and 0.2M NaCl (Figure 6J, blue line), while staircase stress followed a sequential, constant change of 0.2M, without changing back to 0 (Figure 6J, black line). These stresses resulted in differential dynamics for both cell volume reduction and Hog1^nuc^ responses over time (Figures 6K, 6L, S5).

Comparing the observations on Hog1 responses under the five distinct NaCl stress paradigms, namely Linear, Exponential, Sum of Polynomials, Pulsatile, and Staircase Stress, to the quantitative results obtained from different models, confirmed that the Hog1 pathway performs signaling according to a hybrid model (a sum of linear signaling and logarithmic signaling) in response to gradual NaCl stress (Figures 6C, 6F, 6I, 6L compared to Figures 5D, 5H, 5L, 5P, and 5T). These findings collectively confirm that the HOG pathway operates at a signaling mode that incorporates both a linear signaling and a logarithmic signaling mechanism in response to gradual environmental changes. The distinct response patterns observed under different stress conditions highlight the pathway’s ability to perceive and transduce gradual changes in its environment, providing valuable insights into its role in cellular signal transduction.

## DISCUSSION

In this study, we implemented gradual stress profiles and quantified dynamic cellular responses under a wide range of conditions. This general approach is a significant development toward a quantitative investigation of dynamic signaling processes, including mechanistic understanding of cell signaling in physiologic cellular environments. We found that cells grow substantially better under gradually rising stress compared to acute and pulsatile stress (Figures 2 and S1). We then implemented gradual cell stress stimulations and time-lapse microscopy to quantify cell volume change and Hog1 signaling activation dynamics in single cells over time. We found that cell volume reduction depends on the gradual dynamic of the stress (Figures 3 and S2). Interestingly, we uncovered that cell growth rate negatively correlates with cell volume reduction for different gradual stress conditions (Figures 4, S3, and S4). Our results revealed that cells grow optimally upon stress conditions that cause minimal cell volume reduction. Through implementing quantitative models of three different signaling mechanisms, we predicted the signatures of signaling activation dynamics under five distinct stress paradigms, namely linear, exponential, polynomials, pulsatile, and staircase stress (Figure 5). We subsequently compared these model predictions to our experimental measurements, confirming logarithmic signaling contributing to the Hog1 pathway response to sense the relative change in gradual stress over the background concentration over time (Figures 6 and S5).

We find that cell volume reduction, in a stress dynamic-dependent manner, impedes cell growth independent of the rising part of the Hog1 signaling activation, while prolonged signaling under gradual stress may also help cells improve growth. This physiological signaling mechanism is significant because it induces persistent Hog1 activation upon gradual stimulations that maximizes cell survival in severe stress conditions and links cell morphology changes to growth phenotype. These results show that gradual dynamics of extracellular environments impact cell growth and cell volume and our study provides a quantitative framework to investigate what signaling pathways may be involved in these processes.

Our data establishes logarithmic signaling, a signal transduction mechanism of perceiving relative changes of extracellular stress, with a phenotypic consequence in yeast cells for the first time. Such signaling mechanism may be prevalent in pathways such as ERK, Wnt, NF-κB, Tgf-β, PI3K-Akt and may play important roles to regulate distinct cellular responses and functions during important biological processes relevant to human health and disease (57–61).

## ACKNOWLEDGMENTS

GN is supported by NIH R01GM140240, and Vanderbilt Basic Science Dean’s Faculty Fellow Endowed Chair. The authors thank Drs. Brian Munsky and Zachary Fox for their valuable discussions on the manuscript and Dr. Zachary Fox for his comments on the manuscript. This study used resources at the Advanced Computing Center for Research and Education (ACCRE) at Vanderbilt University, Nashville, TN (NIH S10 Shared Instrumentation Grant 1S10OD023680-01 (Meiler).

## AUTHOR CONTRIBUTIONS

Conceptualization, H.J., G.N; Methodology, H.J.; Software, H.J., G.N.; Validation, H.J.; Formal Analysis, H.J.; Investigation, H.J.; Data Curation, H.J.; Writing – Original Draft, H.J.; Writing – Review & Editing, H.J., G.N., Visualization, H.J.; Supervision, G.N., Project Administration, H.J., G.N.; Funding Acquisition, G.N.

## METHODS

### Resource Availability

#### Lead Contact

Further information and requests for resources should be directed to and will be fulfilled by the Lead Contact, Gregor Neuert.

#### Materials Availability

This study did not generate new unique reagents.

#### Experimental model, yeast strain

We employed *Saccharomyces cerevisiae* BY4741 (MATa; his3Δ1; leu2Δ0; met15Δ0; ura3Δ0) as model system. We used a strain in which a yellow-fluorescent protein (YFP) tag was introduced to the C-terminus of the endogenous Hog1 protein in BY4741 cells via homologous DNA recombination (29).

### Method Details

#### 1. Gradual Cell Stimulations Paradigms

To investigate dynamic responses and phenotypes in yeast cells under various gradual stress conditions, we meticulously designed and executed gradual stress profiles spanning a wide range of concentrations and rates. We employed sodium chloride (NaCl, Sigma-Aldrich, S7653) concentration changes based on polynomial and exponential increases over time, ultimately reaching one of the final concentrations of 0.10M, 0.20M, 0.40M, 0.60M, or 0.80M over a 25-minute period in independent experiments (Figures 3C and S3A) (16, 27).

In our experimental setup, we employed programmable syringe pumps (New Era Pump Systems, NE-1200) capable of operating in 251 phases, enabling the creation of profiles comprising 250 linear concentration intervals, each lasting 6 seconds. This design ensured a continuous, monotonically increasing concentration profiles for all conditions in this study. By progressively increasing the rate between intervals, we achieved the desired rising concentration profiles throughout the entire treatment duration. During each 6-second interval, the pump consistently delivered the stress by introducing the appropriate volume of concentrated stress solution, while correcting for both the added and removed volumes. The former step delivered the concentrated stress to a mixing flask, while the latter delivered the changing concentration to the cells over time.

These gradual NaCl concentrations were then applied to yeast cells in real time to evaluate their impact on cell growth, cell volume change, and Hog1 kinase nuclear localization (Hog1^nuc^) phenotypes. Additionally, our experimental setup allowed us to implement step-like profiles and pulsatile and staircase stresses by switching the extracellular medium between different fixed final concentrations of NaCl (16, 27).

#### 2. Concentrated Stress, 5M NaCl in CSM

We prepared a concentrated stress of 5M NaCl in 1X CSM (yeast growth media) for use in all experiments in this study. To create this solution, we dissolved 292.2 grams of sodium chloride (NaCl) in 850 mL of CSM 1X, followed by autoclaving. We then adjusted the final volume to 1,000 mL using 50 mL pipettes and additional CSM. CSM 1X was prepared using 75mL CSM 10X (5.925 g CSM, Formedium DCS0019, in 750 mL ddH2O), 75mL YNB 10X (51.75 g Yeast Nitrogen Base w/o Amino Acids, YNB, Formedium, CYN0410 in 750 mL ddH2O), 75 mL Glucose 20% (150 g Glucose, Fisher, Dextrose Anhydrous 147, M-15722 in 750 mL ddH2O), and 525mL ddH2O (16).

#### 3. Yeast strain and cell culture

We employed *Saccharomyces cerevisiae* BY4741 (MATa; his3Δ1; leu2Δ0; met15Δ0; ura3Δ0) for our experiments. To assess the nuclear enrichment of Hog1 in response to osmotic stress at the single-cell level, we utilized a yellow-fluorescent protein (YFP) tag, which we introduced to the C-terminus of the endogenous Hog1 protein in BY4741 cells via homologous DNA recombination (29).

In preparation for our experiments, yeast cells from a stock stored at -80°C were streaked onto a complete synthetic media plate three days prior to the study. The day before the experiment, a single colony from the CSM plate was selected and used to inoculate 5 mL of CSM medium to establish a pre-culture. After 6-12 hours, we measured the optical density (OD) of the pre-culture and diluted it into fresh CSM medium to achieve an OD of 0.5 the following morning.

#### 4. Cell Growth Measurements

In this study, we exposed yeast cells to various gradual osmotic stress conditions to investigate their effects on cell growth phenotypes. Yeast cell cultures were grown until they reached the exponential growth phase (OD600 = 0.5) before subjecting them to different stress conditions. We monitored cell growth by measuring optical density of cell culture at 600 nm (OD600) over time, both before and after each stress treatment. Our experiments involved repetitive NaCl stress treatments, including long pulses, short pulses, linear increases, and exponential increases along proper controls to account for stress duration, integrated stress exposure, and cell density, with each reaching a final concentration of 3M NaCl. Detailed profiles for each condition can be found in Figure 2 and S1. To implement these conditions, we added a total of 21.75 mL of concentrated stress (5M NaCl in CSM) to 14.5 mL of cells in CSM.

For pulsatile stress treatments, the entire 21.75 mL of 5M NaCl was introduced to the cells in a mixing flask at once. In the case of linear and exponential stress treatments, the 21.75 mL of 5M NaCl was gradually delivered to the cells over time using a 20 mL BD syringe (BD™, 309628) mounted on a syringe pump. Introducing 21.75 mL of 5M NaCl either in a single acute stress event (pulsatile stress) or gradually over 120 minutes (linear and exponential stress) leads to distinct cell dilution profiles thus distinct cell density changes throughout the 2-hour treatment duration (Figure 2A). These variations may have implications for post-stress cell growth. The experiment presented in Figure 2D serves as a control to account for this potential effect. During the gradual stress applications, the syringe pump functioned according to the calculated linear or exponential profile for the delivery of each over the 90-minute treatment period. The dispensed volume selections were determined by the capacity of the 20 mL BD syringes, which allowed for a 21.75 mL fill volume for each experiment without the need for refilling during the treatment.

After each stress treatment, we removed the NaCl by filtering the cells using 0.45 μm filters (Filter Membranes, Millipore, HAWP09000) and a glass vacuum system, and then resuspended them in 40 mL of fresh CSM. We incubated the cells and measured OD600 during the next 5 hours at various time points: 15 minutes, 1 hour, 1 hour and 45 minutes, 2 hours and 30 minutes, 3 hours and 15 minutes, 4 hours, and 4 hours and 45 minutes post-incubation. Throughout the experiments, we ensured homogenous mixing by constantly shaking the cells at 200 rpm in a 30°C incubator, and we maintained the growth media and concentrated stress within the same 30°C incubator for temperature control.

#### 5. Time-lapse microscopy

A volume of 1.5 mL of yeast cells at OD600 = 0.5 were centrifuged, the supernatant was removed, and the remaining cell pellets (approximately 40 µl) from three centrifugation tubes were combined into one, and loaded into the flow chamber. After loading the cells into the flow chamber, the chamber was attached to the microscope, and a 5-minute incubation period allowed the cells to adhere to the Concanavalin (Sigma, L7647) coverslip. The flow chamber configuration comprised an 1/8” Clear Acrylic Slide with three holes (Grace Bio-labs, 44562), a 0.17 µm thick T-shape SecureSeal Adhesive spacer (Grace Bio-labs, 44560, 1 L 44040, R&D), a Microscope Cover Glass (Fisher Scientific, 12545E, No 1 22 × 50 mm), and Micro medical tubing (Scientific Commodities Inc., BB31695-PE/4, 100 feet, 0.72 mm ID × 1.22 mm OD). The cover glass had been coated with a solution of 0.1 mg/ml ConA in H2O.

NaCl stress profiles were established using Syringe Pumps, consistent with the procedures described in previous sections. To facilitate acute stress, flask 1 contained CSM medium with a predetermined NaCl concentration. For gradual stimulations, the mixing flask initially held media without NaCl at time zero, and we generated concentration profiles by pumping 5M NaCl CSM media into the mixing flask using a syringe pump. These media changes were continuously mixed on a magnetic stir plate to achieve the desired profiles to stress the yeast cells.

#### 6. Image Acquisition

For the quantification of nuclear enrichment of Hog1 terminal kinase in individual yeast cells as an indicator of pathway activation, we employed a strain in which a yellow-fluorescent protein (YFP) was integrated at the C-terminus of the endogenous Hog1 protein in *Saccharomyces cerevisiae* BY4741 yeast, using homologous DNA recombination.

Our imaging setup consisted of an inverted microscope, specifically the Nikon Eclipse Ti, which was equipped with several components, including perfect focus (Nikon), a 100× VC DIC lens (Nikon), a YFP fluorescent filter from Semrock, an X-cite fluorescent light source (Excelitas), and an Orca Flash 4v2 CMOS camera from Hamamatsu. The entire system was controlled via the Micro-Manager program.

To capture images, we conducted the following steps:

Utilizing the microscope’s xy-plane movement, we selected a field of view that exhibited an optimal density of yeast cells within a single z-plane. The z-focus was carefully adjusted to visualize the boundary of most cells, appearing as a white ring. Time-lapse images were recorded, encompassing both bright-field images, taken at intervals of every 10 seconds, and fluorescent channel images, acquired at intervals of every minute.

#### 7. Image Segmentation

The published articles (27, 29, 62–64) include the image processing codes for cell segmentation and quantification that are used to analyze the cell volume change and Hog1 nuclear localization data during this study. The codes are available at https://osf.io/kwbe6/.

#### 8. Quantification of Cell Growth rates

To determine cell growth rates, we employed a fitting procedure to the OD600 measurements, from which we calculated the doubling times and growth rates. This involved fitting the OD600 data to doubling functions, calculating doubling times and growth rates, and normalizing these values with respect to the control condition to analyze the impact of various stress conditions on cell growth.

The cell growth data were fitted using the following mathematical function:

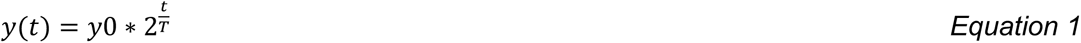

Where:

- *y*(*t*) represents the OD600 measurements at time t.
- *y*0 is the initial OD600 value for each condition.
- *T* represents the doubling time, which is the time it takes for the OD600 to double.
- The reciprocal of *T* represents the cell growth rate.

We independently fitted data from each section for each biological replicate (Figure 2 and S1). For each condition, we performed 30 different fits. The resulting doubling times and growth rate values are obtained as averages of the fit values from these 30 independent fits. To facilitate comparison, we normalized the growth rate values with respect to the mean growth rate observed under control conditions, which involved no NaCl. This normalization allows us to assess the relative changes in growth rates induced by the different stress conditions.

#### 9. Quantification of Cell Volume Change

In this study, the assessment of cell volume changes involved two main quantities:

i. *Relative Cell Volume Changes Over Time:* We quantified the dynamics of cell volume change over time, normalized to the initial timepoints just before the application of each stress condition. This measure allowed us to track how cell volume evolved under different gradual stress.
ii. *Relative Cell Volume Reduction Over Time:* This specific metric was introduced to quantify cell volume change under each stress condition compared to that observed under a no-salt control condition. It was defined as the difference between cell volume changes under a given condition and the baseline cell volume expansion that occurs in standard growth media over time.

To obtain the relative cell volume reduction data, we subtracted the cell volume change under each stress condition from that under the no-salt control condition.

For the cell volume reduction data, we applied polynomial functions to describe the dynamics and compare cell volume reduction to cell stress dynamics. The fitting function took the following form:

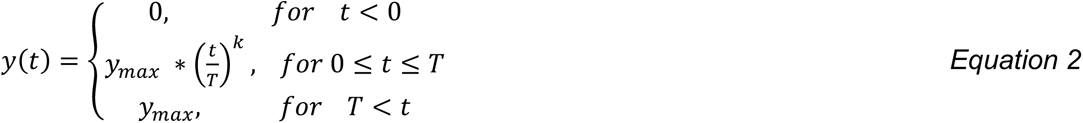

- *y*(*t*) represents the cell volume reduction over time.
- *T* is a fixed value of 25 minutes, as it was the duration of the stress period.
- *y*_&’(_ stands for the maximum volume reduction achieved for each specific condition.
- *k* corresponds to the polynomial order.

We independently fitted the data for each condition between time points 0 and 25 minutes. For each condition, we conducted 30 different fits, and the values of *k* and *y*_&’(_ were determined as averages based on the outcomes of these 30 independent fits. For the results presented in Figure 4C-4G and S3D, we used the following fitting function for the polynomial part:

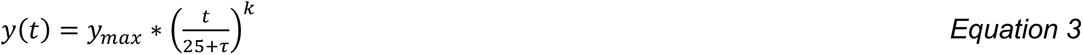

Where, *τ* is a new free parameter that is introduced with its value constrained to the range [-2, 2] in units of minutes, to account for the small delay or expansion that was observed in the measured cell volume data. For example, *τ* = 0 corresponds to the scenario where *y*(*t*) reaches its maximum value of *y*_&’(_ at *t* = 25 minutes.

We examined two key aspects of cell volume change: the maximum cell volume reduction and the total cell volume reduction, both assessed over the 25-minute stress period. This analysis allowed us to quantitatively capture the overall trends in cell volume dynamics under the various stress conditions.

#### 10. Quantification of Hog1 Nuclear Localization

In our study, quantification of Hog1 nuclear localization is a crucial aspect of understanding the signaling dynamics within yeast cells subjected to various gradual stress. This quantification involves the calculation of the ratio of Hog1 signal derived from pixels representing the nucleus of individual cells to the total signal from the entire cell, with corrections made for the image background and photobleaching of the fluorescent signals over time. At the single-cell level and over time, we employ the following formula to quantify Hog1 nuclear localization using the nuclear, total cell, and image background signals over time:

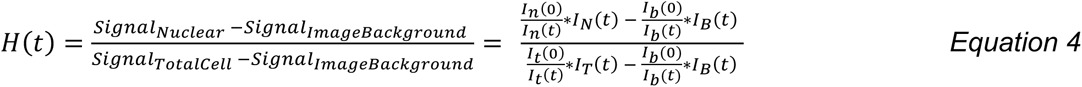

Here, the subscripts (N/n, T/t, B/b) correspond to (nuclear, total cell, image background) regions for a given single cell. We use capital letters (*I_N_*, *I*_T_, *I_B_*) to denote the raw fluorescent signals from these regions. Meanwhile, lowercase letters (*I_n_*(*t*), *I_t_*(*t*), *I_b_*(*t*)) represent the double-exponential decaying function fits for these signals obtained from pre-stress and the end of trajectories, modeling the photobleaching decay, while (*I_n_*(0), *I_t_*(0), *I_b_*(0)) are the values at the first timepoint. It’s important to note that in all conditions we maintain stress at fixed final concentrations for at least 25 minutes allowing Hog1 activities to adapt to pre-stress levels during these fits, allowing for accurate quantification of nuclear localization over time. We quantified Hog1 nuclear localization as Hog1nuc (t) = H(t) – H(0), where H(0) is the value at pre-stress timepoint.

This analysis plays a crucial role in correcting for the decay in fluorescent signal attributable to photobleaching, a phenomenon that can impact the accuracy of Hog1 nuclear localization measurements. For our control condition (CSM growth media with 0M NaCl), the method effectively achieves a constant Hog1nuc(t) value of 0 over time. However, the quantification of nuclear localization is influenced by the photobleaching events associated with the fluorescent protein used to label Hog1.

To address this issue and ensure accurate quantification, we conducted additional control experiments involving step-like stimulations applied to the cells at two distinct time points, namely t=12 minutes and t=20 minutes (Figure S2A-S2C). These experiments resulted in an additional 12 and 20 fluorescent images captured before the change in stress, compared to experiments in which we applied the step stress at t=0. The Hog1nuc(t) values obtained from these experiments, using the aforementioned method, exhibited variations in signaling levels.

In response to these observed variations, we undertook supplementary experiments to empirically correct for the impact of photobleaching in Hog1nuc quantification. Specifically, we conducted nine experiments involving a step-like change of NaCl concentration to 0.6M at t=0, each with an increasing number of pre-stress photobleaching snapshots taken (Figure S2D-S2G). This resulted in varying levels of Hog1nuc, depending on the number of pre-stress photobleaching snapshots.

To quantify these outcomes, we employed a fitting procedure to establish a relationship between the Hog1nuc values and the number of pre-stress photobleaching snapshots. Importantly, all our experiments involved the acquisition of fluorescent signal data at one-minute intervals, providing us with a quantitative factor to correct Hog1nuc signal over time (Figures S2H-S2J). This correction procedure ensures that the impact of photobleaching on Hog1 nuclear localization is appropriately accounted for in our analyses.

Similar to cell volume reduction data, we applied polynomial functions to describe the rising dynamics of Hog1^nuc^ data over time and compare them to that of cell volume reduction and cell stimulations. The fitting function is given by Equation 2, where in these fits, we fix T to the value of time (in units of minutes) at which Hog1^nuc^ reaches its maximum value. The results from these analyses are presented in Figures 4 and S4.

#### 11. Quantitative Modeling of Different Signaling Mechanisms

To establish a relationship between time-dependent stress and the resulting signaling response dynamics, we developed a series of quantitative models to investigate various signaling mechanisms. Our goal was to predict the signaling response to different gradual stress accurately. To conduct a comprehensive analysis, we compared the behaviors of the following models:

1. *Linear signaling:* This model is based on rate sensing, where the cellular response is directly proportional to the rate of change in the stress over time. In this model, we describe the signaling response as the first time-derivative of the stress profile.
2. *Logarithmic signaling:* This model is based on relative rate sensing, in which the response is proportional to the relative change in stress concerning the background concentration over time. In this model, we describe the signaling response as the ratio of the first time-derivative of the stress divided by the stress concentration over time.

We considered various gradual stress functions, as shown in Figure 5, and quantified the response from each model using each of the three approaches. Specifically, we numerically simulated concentration values at pre-specified timepoints and from those values we calculated the rate of change as difference in concentration values between adjacent timepoints. We used these values under gradual stress profiles to calculate responses for each model as described above. We then applied a 3-point moving average to smooth the responses.

### Statistics on Single Cells

### Additional resources

*For complete details on the protocol, please refer to* https://star-protocols.cell.com/protocols/786.

## SUPPLEMENTAL INFORMATION

**Figure S1.**
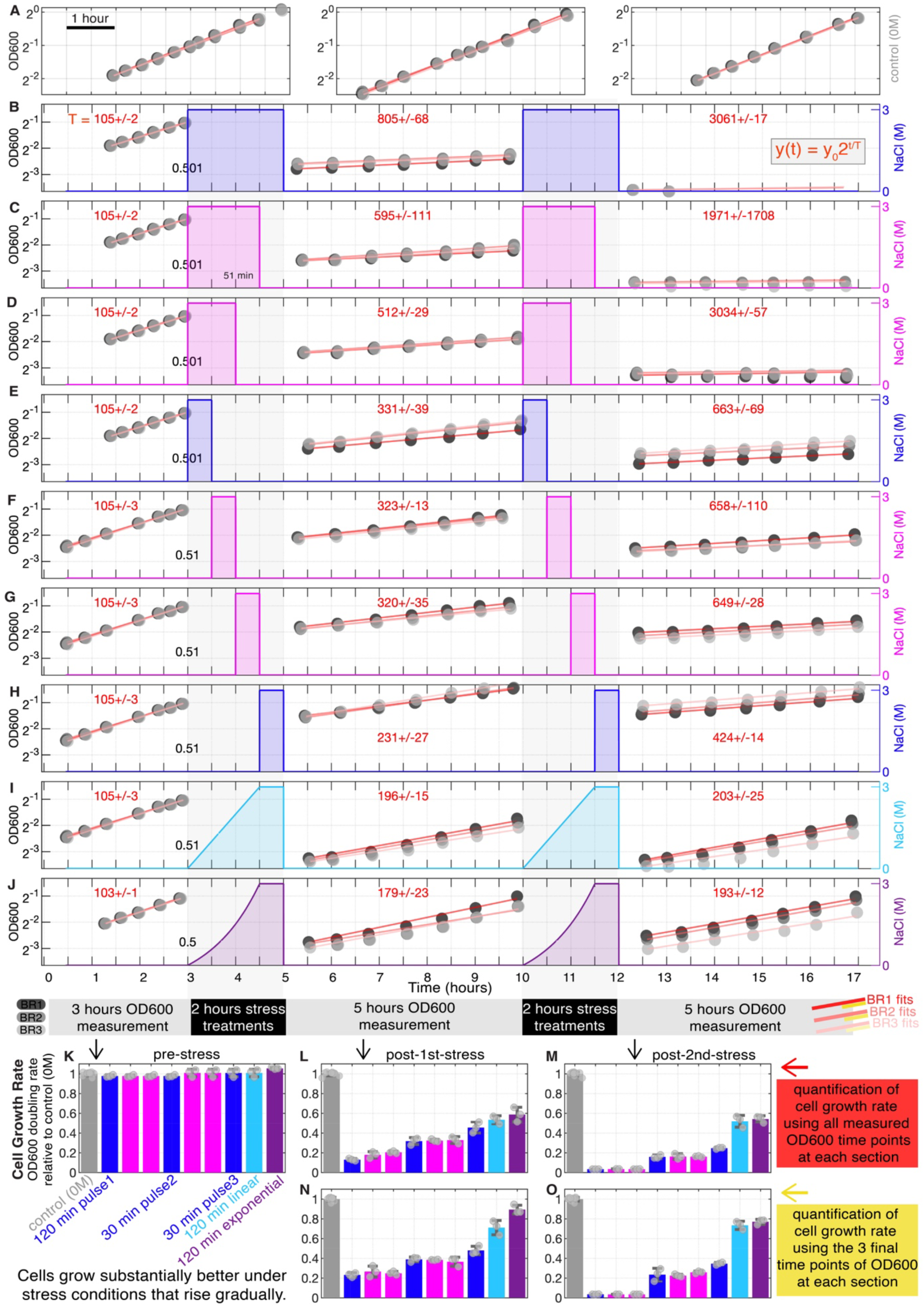
Cell Growth Phenotype under Various Gradual NaCl Stress Conditions. (**A**-**O**) Cell growth phenotypes were quantified over time under different NaCl stress conditions using OD600 measurements of bulk yeast cell cultures. (**A**) OD600 measurements for yeast cells growing in normal CSM media without NaCl addition over time in three independent experiments, each with three independent biological replicates. Black, gray, and light-filled circles represent the measurements, and red lines show the fits to an exponential function in the log2 y-axis plot over time. (**B**) Cells were grown to the exponential phase (OD600 = 0.5) at the start of each treatment. Subsequently, they were exposed to various NaCl treatment profiles, including (**B**-**E**) repetitive 3M NaCl osmotic stresses in the form of pulses with durations of (**B**) 120 minutes, (**C**) 90 minutes, (**D**) 60 minutes, and (**E**) 30 minutes, each applied to the cells at t=0 of the 2-hour treatment period. (**F**-**H**) Three 30-minute pulse NaCl treatments were administered to the cells at (**F**) t=30 minutes (**G**) t=60 minutes, and (**H**) t=90 minutes from the start of the 2-hour treatment period. We also considered (**I**) a 120-minute linear (cyan) and (**J**) a 120-minute exponential (purple) increase. Each linear and exponential stress profile gradually increased to 3M in 90 minutes and remained at 3M for another 30 minutes. At the end of each 2-hour treatment period, cells were filtered to remove NaCl, re-suspended in fresh cell growth media, and their OD600 values were monitored for 5 hours in three biological replicates (represented by black, gray, and light-filled circles in **A**-**J**). After 5 hours, another round of 2-hour stress treatments (following the protocols described in **B**-**J**) and 5-hour OD600 measurements were repeated. During each pre-stress, post-1^st^-stress, and post-2^nd^-stress period, OD600 measurements from each BR were fitted to an exponential growth curve (**A**-**J)**, fits are shown in red in a log2 y-axis plot over time. Cell growth doubling times were calculated from each fit and are shown in red numbers rounded to the nearest minute. (**K**-**O**) Cell growth rates following each stress treatment were quantified using OD600 doubling rates relative to the control (0M, no stress) at (**K**) pre-stress, (**L, N**) post 1st stress treatments, and (**M, O**) post 2nd stress treatments. Results in (**L**, **M**) are based on fitting all timepoints, while the results in (**N**, **O**) are based on fitting only the measurements from the three final timepoints.

**Figure S2.**
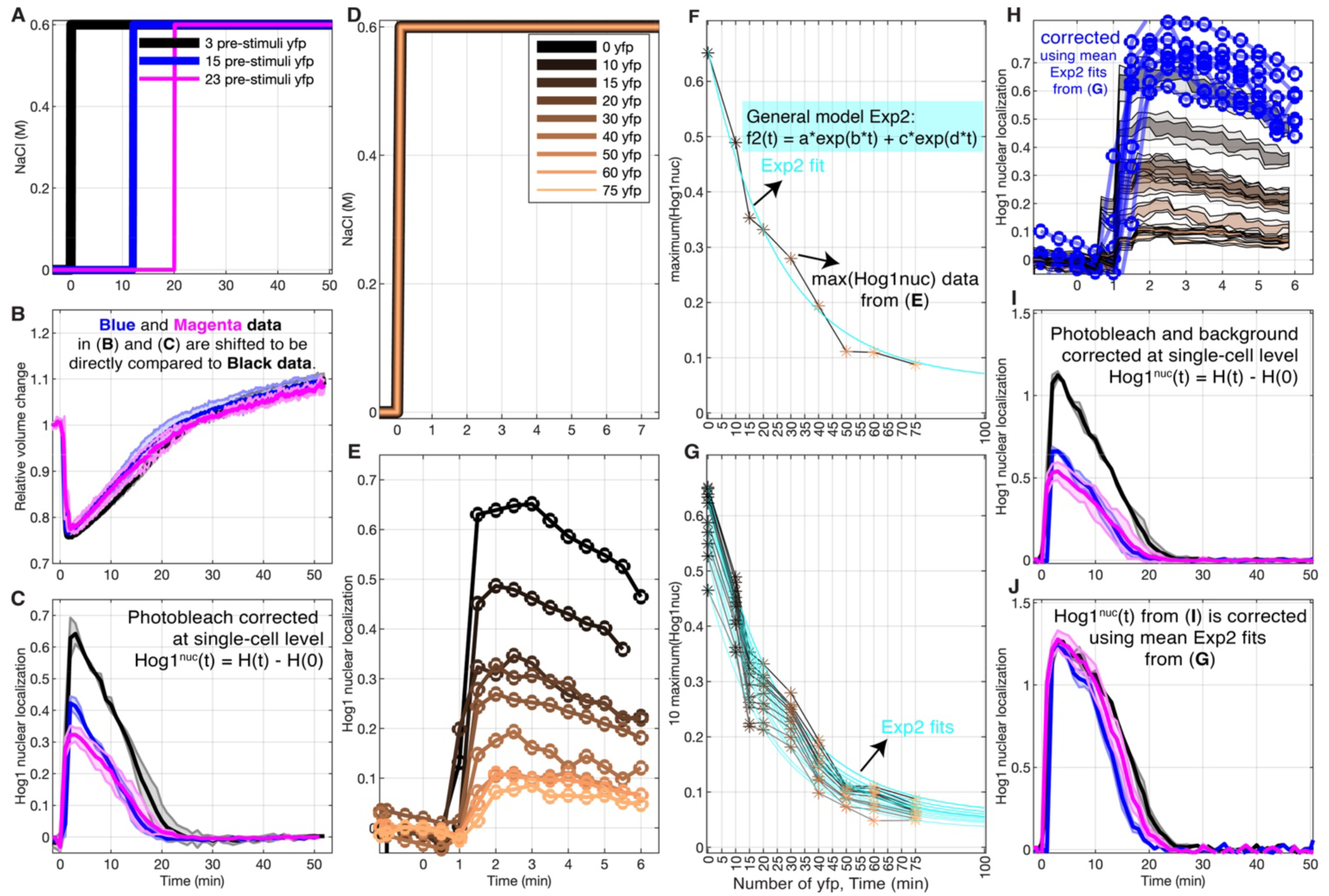
Correction for Photobleaching of Hog1 Nuclear Localization Data. (**A**) Three step-like NaCl stress with final concentrations of C_max_ = 0.6M were applied to yeast cells at t = 0 minutes, 12 minutes, and 20 minutes in three independent experiments. Images were captured in brightfield every 10 seconds and fluorescent images every one minute with exposure times of 10 and 20 milliseconds (see Methods). The number of fluorescent images (yfp) captured before the step change in stress varied in these experiments, including 3, 15, and 23 fluorescent images, respectively. These stress resulted in (**B**) consistent relative cell volume changes, which were quantified using brightfield images. However, they led to (**C**) varying Hog1 nuclear localization dynamics (Hog1nuc) due to photobleaching of fluorescent images over time. (**D**) To address this issue empirically, we conducted additional control experiments involving step-like 0.6M NaCl stress applied to the cells all at t=0, each with an increasing number of pre-stress photobleaching snapshots taken, as shown in the legend. This resulted in (**E**) varying levels of Hog1nuc, depending on the number of pre-stress photobleaching snapshots. (**F**) The maximum Hog1nuc values were quantified as a function of the number of pre-stress photobleaching snapshots and fitted to a two-term exponential model to calculate the decay rate per yfp image. Fluorescent images were captured every one minute in all experiments, allowing quantification of photobleaching effects in Hog1nuc data over time. (**G**) The 10 highest Hog1nuc values were quantified as a function of the number of pre-stress yfp snapshots and fitted with a two-term exponential model to calculate the decay rate per yfp image. The decay term calculated as an average from these 10 Exp2 fits was then used to correct all the Hog1nuc data, as follows: corrected_Hog1nuc(t) = (f2(0)/f2(t)) * Hog1nuc(t), where f2(0) represents the value of f2(t) at t=0. (**H**) The corrected Hog1nuc data from (**E**) are shown in blue, and the copper colormap in (**H**) corresponds to data in (**E**). The outer shaded plots represent the 40-60 percentile of single cells for each condition. (**I**) Hog1nuc was quantified under stress in (**A**), correcting both photobleaching of fluorescent images over time and background fluorescent intensity at single cells as described in the Methods section using Equations 4 in the Methods section. Y-axis represent per pixel nucleus to per pixel total cell fluorescent intensity. (**J**) The Hog1nuc data from (**I**) were corrected using our empirical method described in (**G**).

**Figure S3.**
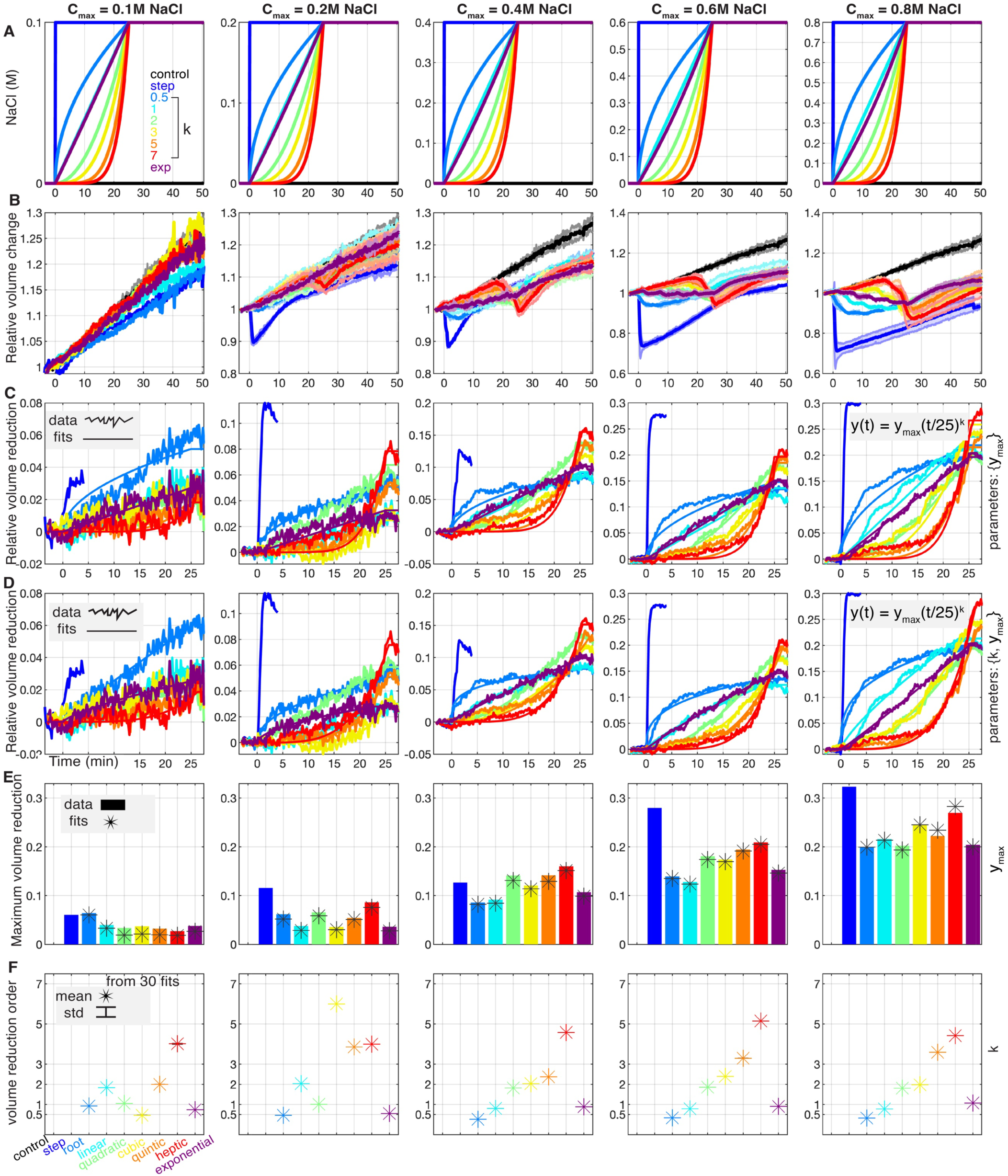
Dynamics of Cell Volume Reduction in Response to Gradual NaCl Stress. (**A**) Yeast cells were subjected to gradual NaCl stress, with final concentrations ranging from C_max_ = 0.1M to 0.8M, resulting in differential (**B**) relative cell volume changes and (C-D) cell volume reduction phenotypes over time. The thick lines and shaded areas in (**B**) represent the mean and standard deviation (std) from 2 or 3 biological replicates, each consisting of approximately 100 single cells observed through live cell time-lapse microscopy. In (**C-D**), the noisy lines represent the data (averages from biological replicates), while the smooth lines represent the fits of the data to polynomial functions. The fits involve one free parameter {y_max_} in (**C**) and two free parameters {y_max_ and k} in (**D**). Further details on the fitting procedure can be found in the Methods section. For each condition, the values of k and y_max_ were determined as averages based on the outcomes of 30 independent fits and are presented in (**E-F**), along with their values directly quantified from the data. (**E**) The maximum cell volume reduction over the 25-minute stress period exhibited variability across different gradual stresses, regardless of the NaCl stress final concentration. The fit values for y_max_ here are from (**D**). (**F**) The fit values for the polynomial order, k, are shown, obtained from the fits in (**D**).

**Figure S4.**
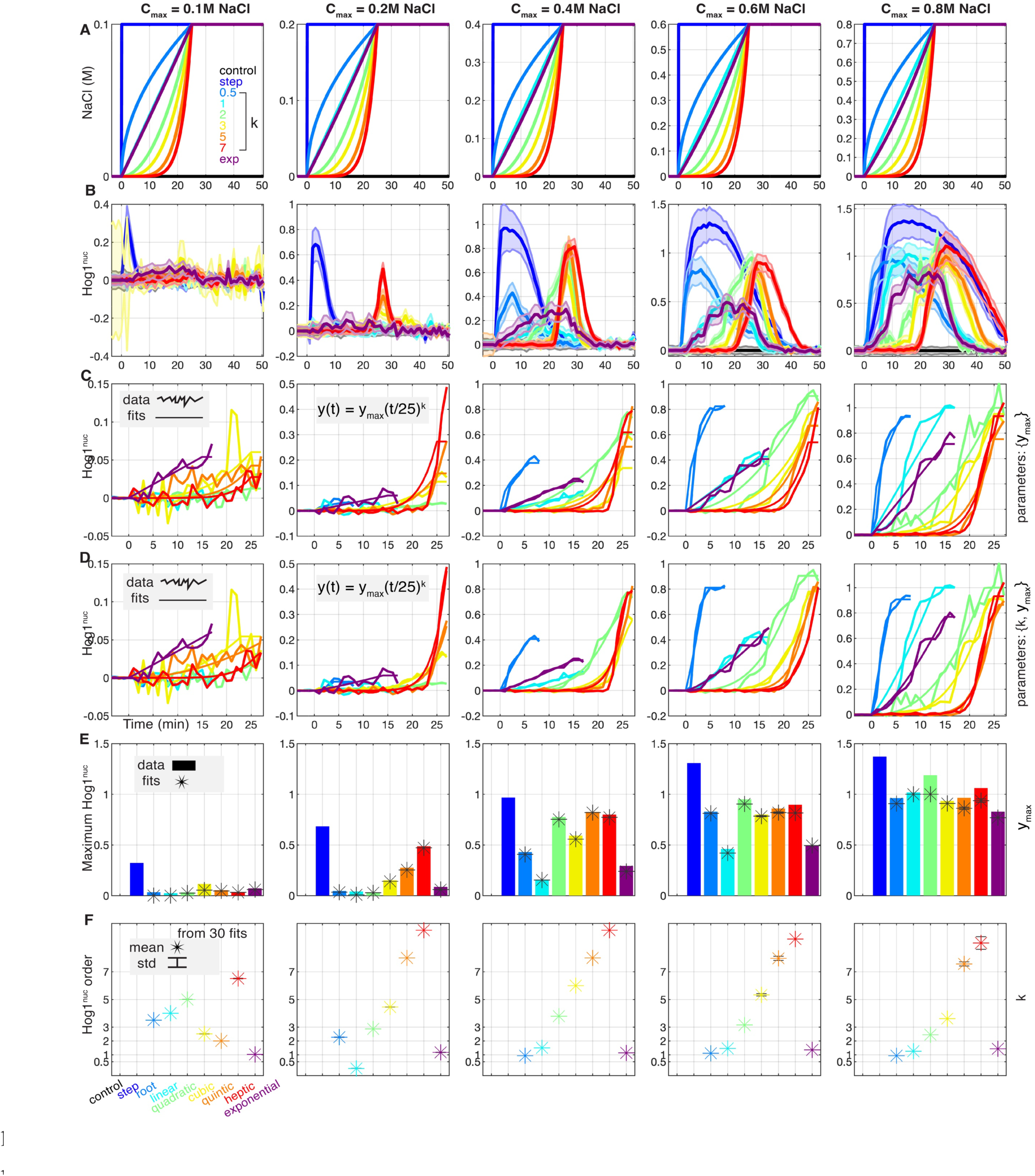
Dynamics of Hog1 nuclear localization in Response to Gradual NaCl Stress. (**A**) Yeast cells were subjected to gradual NaCl stress, with final concentrations ranging from C_max_ = 0.1M to 0.8M, resulting in differential (**B-D**) Hog1 nuclear localization over time. The thick lines and shaded areas in (**B**) represent the mean and standard deviation (std) from 2 or 3 biological replicates, each consisting of approximately 100 single cells observed through live cell time-lapse microscopy. In (**C-D**) we depict the ascending Hog1nuc data to examine the activation dynamics from the stress change until Hog1 reaches its maximum value. The comprehensive analysis of Hog1 activation following its peak or early rise is presented and analyzed in Figures 5 and 6. Here, the noisy lines represent the data (averages from biological replicates), while the smooth lines represent the fits of the data to polynomial functions. The fits involve one free parameter {y_max_} in (**C**) and two free parameters {_ymax_ and k} in (**D**). Further details on the fitting procedure can be found in the Methods section. For each condition, the values of k and y_max_ were determined as averages based on the outcomes of 30 independent fits and are presented in (**E-F**), along with their values directly quantified from the data. (**E**) The maximum Hog1nuc over the 25-minute stress period exhibited variability across different gradual stresses, regardless of the NaCl stress final concentration. The fit values for y_max_ here are from (**D**). (**F**) The fit values for the polynomial order, k, are shown, obtained from the fit in (**D**). These k values along with the values from Figure S3F are used to directly compare the dynamics of cell volume change to Hog1nuc dynamics as presented in Figure 4G.

**Figure S5.**
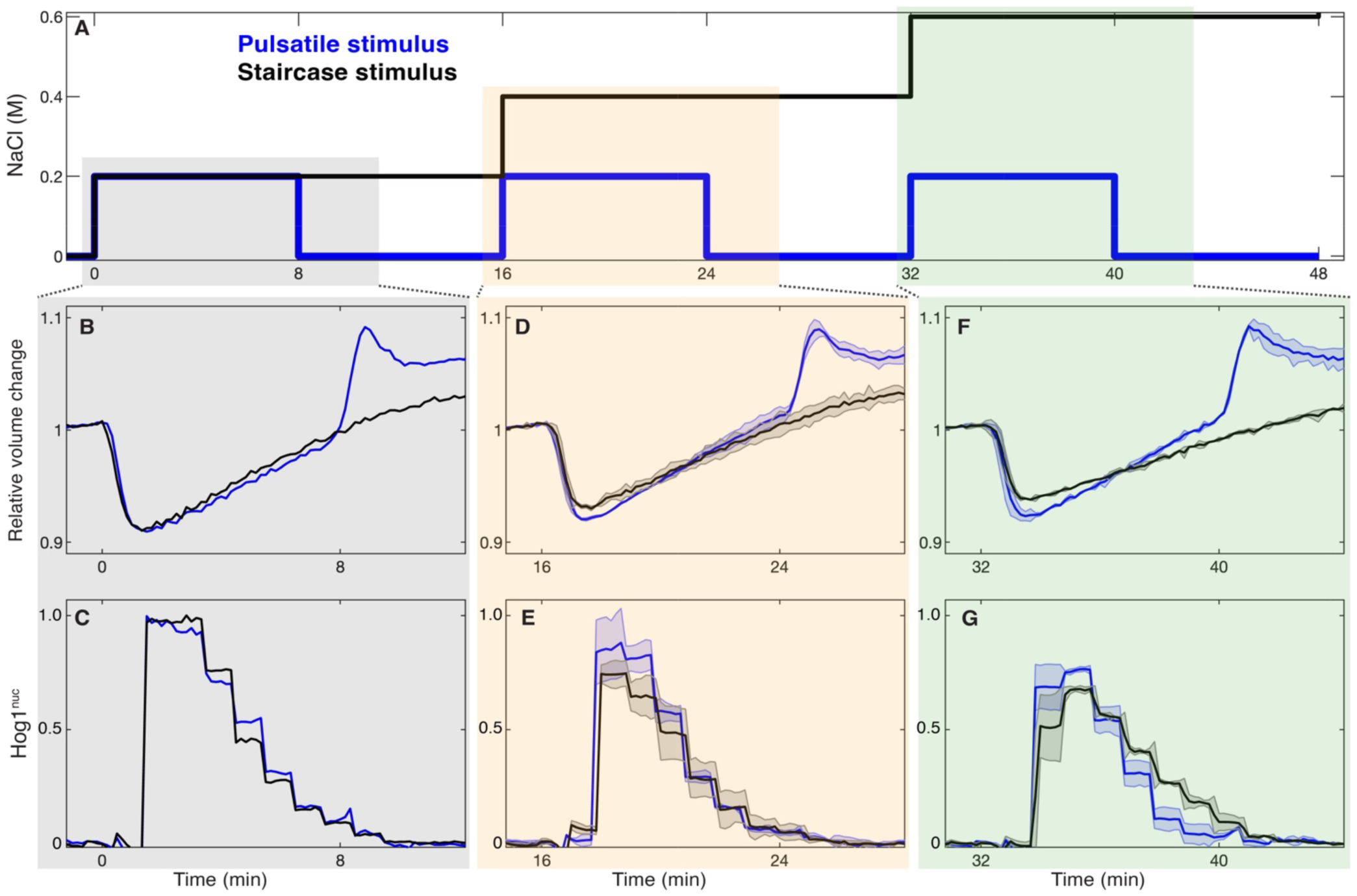
Responses to Pulsatile Versus Staircase Stress. Yeast cells were subjected to gradual NaCl stress in the form of (**A**) pulsatile versus staircase stress, each comprising three periods with identical 0.2M step increases at the beginning of each period, with each period lasting 16 minutes. Relative cell volume changes and Hog1 nuclear localization over time are shown for the (**B, C**) first stress period, (**D, E**) second stress period, and (**F, G**) third stress period, respectively. Imaging was conducted independently at each stress period in separate experiments to minimize the photobleaching effect. The lines in (**B, C**) represent the means from hundreds of single cells. The thick lines and shaded areas in (**D-G**) represent the means and standard deviations (std) from 2 biological replicates, each consisting of approximately 100 single cells observed through live cell time-lapse microscopy.

